# Efficient Sequencing, Assembly, and Annotation of Human KIR Haplotypes

**DOI:** 10.1101/2020.07.12.199570

**Authors:** David Roe, Jonathan Williams, Keyton Ivery, Jenny Brouckaert, Nick Downey, Chad Locklear, Rui Kuang, Martin Maiers

## Abstract

The homology, recombination, variation, and repetitive elements in the natural killer-cell immunoglobulin-like receptor (KIR) region has made full haplotype DNA interpretation impossible without physical separation of chromosomes. Here, we present a new approach using long-read sequencing to efficiently capture, sequence, and assemble diploid human KIR haplotypes. Sequences for capture probe design were derived from public full-length gene and haplotype sequences. IDT xGen® Lockdown probes were used to capture 2-8 kb of sheared DNA fragments followed by sequencing on a PacBio Sequel. The sequences were error corrected, binned, and then assembled using the Canu assembler. The assembly was evaluated on 16 individuals (8 African American and 8 Europeans) from whom ground truth was known via long-range sequencing on fosmid-isolated chromosomes. Using only 18 capture probes, the results show that the assemblies cover 97% of the GenBank reference, are 99.97% concordant, and it takes only 1.8 contigs to cover 75% of the reference. We also report the first assembly of diploid KIR haplotypes from long-read WGS, including the first sequencing of cB05∼tB01, which pairs a *KIR2DS2*/*KIR2DS3* fusion with the tB01 region. Our targeted hybridization probe capture and sequencing approach is the first of its kind to fully sequence and phase all diploid human KIR haplotypes, and it is efficient enough for population-scale studies and clinical use.

## Introduction

The 9 protein coding killer-cell immunoglobulin-like receptor (KIR) genes occur in 12 loci, span ∼10-16 kb each, and alleles from any two genes are over 85-98% identical. Frequent recombination throughout the 100-200 kb haplotypes has made their order and copy number highly variable. The genes encode antigen-recognition proteins that initiate signaling pathways in natural killer (NK) cells. The proteins recognize human leukocyte antigen (HLA) and its peptide, whose recognition can lead to the death of the target cell (infected, cancerous, foreign, etc.). NKs and their KIR receptors are essential to human health and have functional roles that impact viral infections, pregnancy, autoimmune diseases, transplantation, and immunotherapy(1)(2)(3)(4)(5)(6)(7).

Genetic interpretation of exonic-or-lower resolutions from next generation sequencing (NGS) is often ambiguous and unphased, and therefore limits precise understanding of how KIR sequences affect phenotypes. The importance of high-throughput high-resolution typing is exemplified by the fact that the genes contain extensive exonic SNP and short insertion/deletion (indel) variations which rivals that of its binding partner: HLA class I(1)(8). Over 300 full-length DNA and almost 1000 protein reference alleles have been reported in IPD-KIR(9). All resolutions except haplotyping are ambiguous or require statistical phasing from few references. This may be an important issue, since KIR genes are homologous, compact, and regularly interspersed with related LINE1 and Alu transposons and they may be regulated in a coordinated fashion(10). A cost-effective high-throughput method that could characterize *all* the sequences within the KIR haplotypes *in cis* could advance that understanding and clarify previously ambiguous and/or contradictory evidence. To date, the only approach for full haplotyping was to physically separate the chromosomes for subsequent sequencing(11)(12)(13)(14)(15), a process whose expense has generally prohibited its use in large-scale association studies. While high-resolution haplotyping by fosmid clones or full gene by PCR is costly and inefficient, low resolution genotyping of gene presence/absence or copy number provide limited information for functional analysis and association tests.

Like much of chromosome 19, the KIR region is dense with repetitive elements, which have provided the mechanisms for its recent evolution by tandem duplication and homologous recombination events. Dozens of distinct gene-content haplotypes are seen in Europeans alone(12)(16)(13)(17). Previous reports have documented over ten distinct common haplotype structures(18). Figure 1 provides an overview of the most common haplotype structures and their informal names. KIR haplotypes are named in two halves: ‘c’ for centromeric (i.e., proximal) and ‘t’ for telomeric (i.e., distal) separated by a recombination hotspot contained in the ∼10 kb intergenic region between *KIR3DP1* and *KIR2DL4*. Each half is also labelled ‘A’ or ‘B’, designating one of two families of haplotypes, based on the gene content(19). The A family haplotype denotes haplotypes with one main gene content structure and relatively large allelic variation. The B family of haplotypes denotes a class of haplotypes with relatively more structural variation and less allelic variation. The haplotype named ‘cA01∼tB01’, for example, means the first (01) centromeric A region *in cis* (‘∼’) with the first telomeric B region.

**Figure 1.**
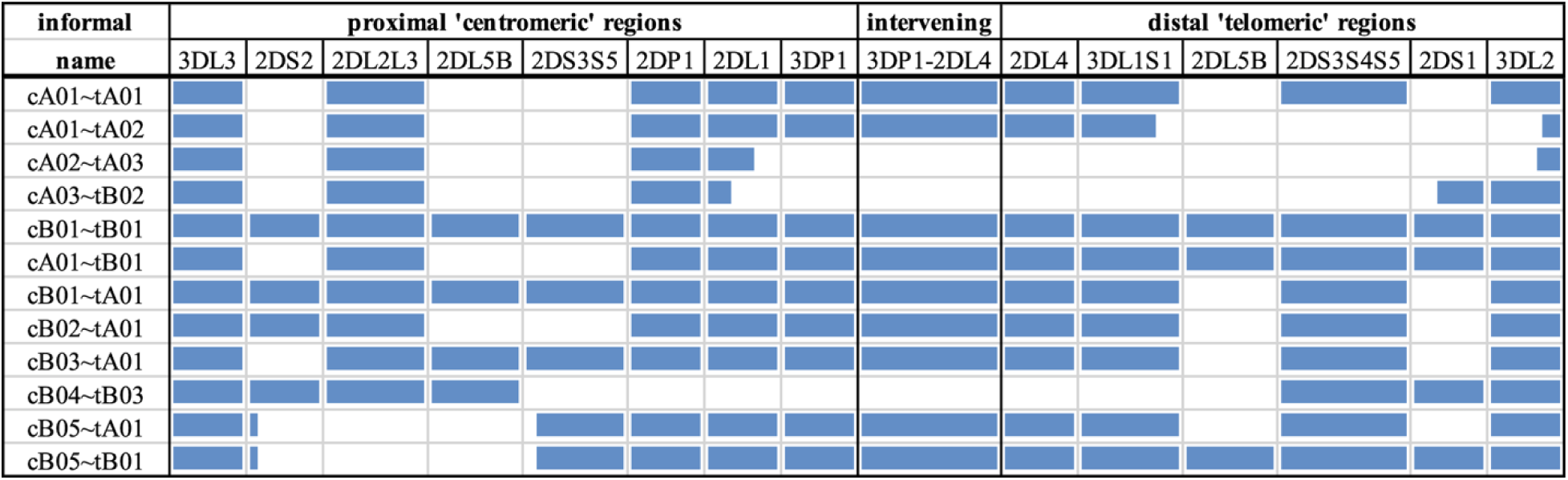
Common KIR haplotype structures and their names. Blue regions represent the presence of a gene or the *KIR3DP1-KIR2DL4* intervening intergenic region. Some genes are partially blue to indicate their portion of a fusion allele. The first column contains the informal name for the haplotype. Each haplotype name is a combination of its two regions: centromeric (proximal) regions are preceded with a ‘c’ and telomeric (distal) regions are preceded with a ‘t’. Not all haplotype structures are shown.

It is difficult to interpret KIR haplotypes for an individual human genome given the reads from high-throughput sequencing when the structural arrangements are unknown. This is largely due to read lengths from prevailing technologies being too short to map unambiguously to the repetitive and homologous KIR genes. Even if the reads could be uniquely placed, they require statistical phasing that is difficult due to lack of phased high-resolution reference libraries. As a consequence, the reads from the KIR region are largely ignored, mis-interpreted, or under-interpreted in current whole genome sequencing (WGS) studies. Therefore, the properties of the KIR region require more careful and specific interpretations than most other regions in human genome.

Here we present an approach that leverages PacBio’s long-read circular consensus sequence (CCS) reads to span DNA homology, and gene homology to efficiently capture 2-8 kb fragments of DNA. It is a workflow to capture, sequence, assemble, and annotate diploid human KIR haplotypes. And it also has broader implications to other genomic regions with variable or repetitive regions alternating with constant regions. When applied to a cohort of 8 African Americans and a cohort of 8 Europeans, the results demonstrate that every KIR gene and intergene contains constant regions that are targetable by capture probes, and that by targeting the constant regions, the variable regions can be captured and sequenced by standard PacBio workflows. Further, maximizing this paradigm shows that 18 short probe sequences can capture KIR haplotypes and allow their unambiguous assembly. Finally, this is an efficient approach that requires no prior knowledge of the individual or references, only utilizes standard lab workflows, and is available in free and open software.

## Materials and Methods

### Overview

The goal of the experiment was to create a set of capture probes and a bioinformatics workflow to efficiently assemble full KIR haplotypes from PacBio CCS reads. The experiments to capture, sequence, assemble, and annotate are depicted graphically in Figure 2. The major steps consist of

**Figure 2.**
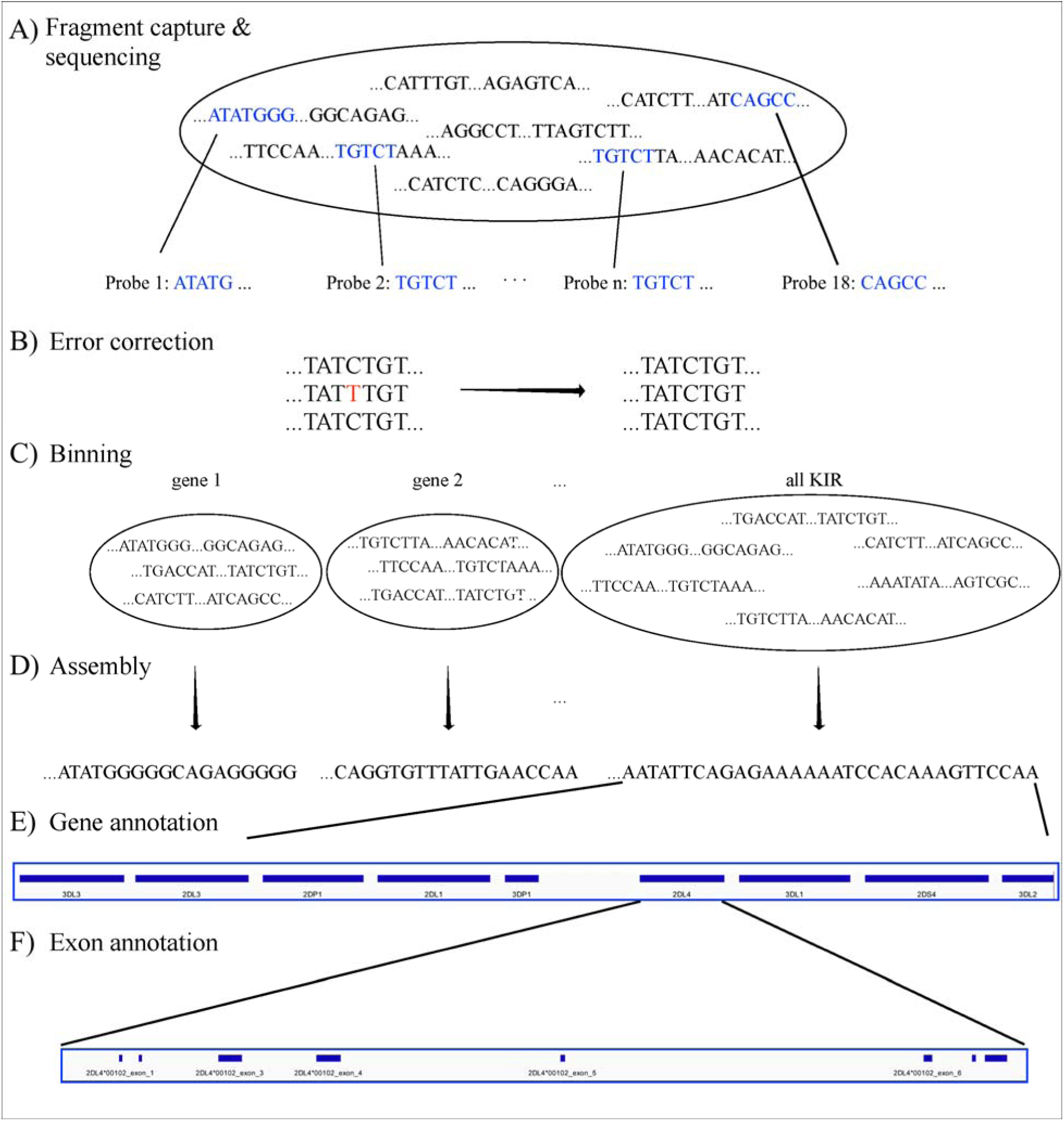
Workflow. The workflow starts with fragment capture *in vitro* or *in silico* and PacBio assembly (A). The sequences are error corrected (B) and binned by KIR region and KIR gene (C) before de novo assembly of each bin (D). Finally, the assembled contigs/haplotypes are annotated by gene (E) and exon (F).

1. Design capture probes.
2. Use the probes to capture the KIR DNA fragments *in vitro* or *in silico* per individual.
3. Sequence the fragments on PacBio Sequel.
4. Error correct the sequences.
5. Bin the sequences per KIR region and gene.
6. *de novo* assemble all the sequences together and each gene bin separately.
7. Annotate the assembled contigs.

### Step 1: Design capture probes

Published KIR haplotypes sequences (36 at the time of this study) as well as all allele sequences from IPD-KIR 2.7.1(20) were used to generate 200 candidate capture probes. The design of the 120-base probes was coordinated with a combination of automated and manual Integrated DNA Technologies (IDT) design tools, including a strain typer alignment tool and a xGen® Lockdown probe design tool. The candidate set of probes was reduced by leveraging sequence homology. First, the haplotype sequences were aligned to two KIR haplotypes (GenBank accessions GU182358 and GU182339). These two reference haplotypes, which together contain all KIR genes, were annotated via RepeatMasker(21)(22). The candidates were prioritized by the highest number of times each aligned to the two reference sequences but not to repetitive elements. The set was chosen by iteratively adding the probe with the highest alignment hit count until both reference haplotypes were covered by less than the expected average DNA fragment length of ∼4-5 kb. Ultimately, experiments were conducted on the original set of 200, a minimal set of 15, and a refined set of 18 capture probes.

### Steps 2-3: Capture and sequence the DNA fragments per individual

Targeted hybridization probe capture and sequencing was performed as previously described(23) with the following modifications (Figure 2A). Human Genomic DNA was sheered with Covaris G-tubes to 6-8 kb followed by 1:1 PB AMPure bead cleanup. Individual specimens were then library prepped with the KAPA library prep kit (Roche) which consisted of end repair and ligation of uniquely barcoded adapters that also contained the PacBio Universal Primer sequence. Samples were enriched with eight PCR cycles using LA Taq (Takara) and PacBio Universal primers followed by a 1:2 PB AMPure bead cleanup. Sample concentrations were measured on the Promega Quantus and 2 μg was size selected for greater than 2 kb fragments on the Blue Pippin System (Sage Sciences).

Eight multiplexed samples for Sequel sequencing were pooled at this point at 0.25 μg per sample. Size selected DNA was then targeted for hybridization probe capture with IDT xGen® lockdown probes according to manufactures instructions with 2 μg of COT DNA. After capture, DNA was removed from streptavidin beads and further enriched with fifteen PCR cycles using LA Taq. DNA fragments were then library prepped and sequenced eight samples per SMRT cell on the Sequel, according to PacBio instructions. Raw sequence data was then demultiplexed (if necessary) and CCS reads generated on SmrtLink 6.0 using 99.9 % subread accuracy filter for generation.

### Steps 4-6: Correct, bin, and assemble the sequences

For both targeted and WGS, the fastq sequences were error corrected with LoRMA(24). The sequences were binned for on-KIR and also binned per genic or intergenic region in silico (Figure 2B-C) using the 18 capture probes for on-off KIR detection and 32,230 gene probes. The gene probes are 25mers and are detailed in a recent manuscript by Roe et al. that has been submitted for peer review and preprinted on bioRxiv(25). Synthetic probe matching was conducted via bbduk(26) with parameters ‘k=25 maskmiddle=f overwrite=t rename=t nzo=t rcomp=t ignorebadquality=t’. This effectively removed any off-KIR sequences and binned the sequences into 15 loci: 14 genes and the intergenic region between *KIR3DP1* and *KIR2DL4*. Sequences in each bin were *de novo* assembled with Canu 2.0(27) (with default parameters except ‘genomeSize=200k’) for each bin separately and all KIR sequences together (Figure 2D). The assemblies utilized only the captured sequences and were not assisted by any prior information, including individual genotypes or reference libraries.

### Step 7: Annotate the assembled contigs

The capture probes were aligned to the contigs, and their patterns allowed gene-specific sequences to be extracted from the haplotypes (Figure 2E). The details are presented in a recent manuscript by Roe et al. that has been submitted for peer review, but at a high level, the algorithm uses the bowtie2 alignment pattern of the 18 capture probes across the contigs/haplotypes to define locus-specific features. Within each feature, the locations of the exons, introns, and untranslated elements were located by searching for inter-element boundaries with 16 base sequences as defined by the full haplotype MSA of 68 human haplotypes (Figure 2F). Each sequence contains 8 bases from one region and 8 from the other. For example, ACACGTGGGTGAGTCC spans the boundary between *KIR2DL4*’s exon two and intron two; the first eight characters are from the second exon, and the last eight bases from the second intron. Almost all boundaries are defined by one or two sequences. The locations of these elements allows for the annotation of protein, cDNA, and full-gene alleles with respect to names assigned in IPD-KIR. The contigs were ultimately annotated in GenBank’s gbf and tbl formats. BioJava(28) was used for some of the sequence processing. Reports on the assembly (or the raw sequences) were generated from Minimap2(29), Qualimap(30), NanoPack(31), QUAST(32), Simple Synteny(33), and Tablet(34).

### Evaluation of the workflow

The capture-sequence-assemble workflow was evaluated on a cohort of 16 individuals whose haplotypes had previously been sequenced using fosmid separation and long-read sequencing(14)(35). Eight are of European (EUR) ancestries (GenBank haplotype sequences KP420437-9, KP420440-6, KU645195-8, and KU842452) and eight of African American (AFA) ancestries (GenBank haplotype sequences MN167507, MN167510, MN167512, MN167513, MN167518, MN167519, and MN167520-9). The European haplotypes are detailed in Roe et al. 2017(18), and the African Americans are detailed in the previously mentioned manuscript by Roe et al. that has been submitted for peer review. The distribution of haplotype structures in the European cohort is 8 cA01∼tA1, and 1 each of cA01∼tB01, cA01∼tB04, cA02∼tA03, cA03∼tB02, cB01∼tB01, cB02∼tA01, and cB04∼tB03; one individual is homozygous for cA01∼tA01 to within a few variants. The distribution of African American haplotypes is 5 cA01∼tA01, 3 cB01∼tA01, 3 cB03∼tA01, 2 cB01∼tB01, 1 cA01∼tA02, 1 cA01∼tB01, and 1 cB02∼tA01. Further details are provided in Supplementary Table 1.

### WGS

Theoretically, if KIR reads could be removed from WGS, the workflow should be able to assemble haplotypes the same as from targeted sequences. To test this hypothesis, whole genome CCS reads were obtained for an Ashkenazim individual (isolate NA24385) from the Genome In a Bottle (GIAB) consortium, as described in Wenger et al. 2019(36). KIR ground truth was unknown previously. KIR reads were separated from WGS as described above and from there the workflow proceeded as usual from the error correcting step (Figure 2B).

## Results

Assemblies were evaluated with ground truth in 16 individuals (32 haplotypes) comprising 11 distinct haplotype structures. Figure 3 depicts the 11 structures as connections between the same genes in different haplotypes and shows how the structures represent expansion and contraction of the A and B haplotype categories across the *KIR3DP1-KIR2DL4* hotspot. Table 1 shows the results of the assembly compared with the reference sequences. For the 8 Europeans on average, the full set of 200 candidate probes provided 98% coverage, with 99.98% concordance, and it took 1.1 contigs to cover 75% of the reference (LG75). When a set of capture probes was reduced to a select 15 and evaluated on both cohorts, the European coverage lowered to 93%, with the same concordance rate (99.98%), and 1.3 LG75. The results for African Americans were very similar: 92% coverage, 99.98% concordance, and 1.6 LG75. When a select 3 more probes were added for a total of 18 capture probes, the assemblies for the 8 African Americans improved to 97% coverage, with 99.97% concordance, and 1.8 LG75. Most of the missing coverage occurred at the 3’ end of the haplotypes: in certain *KIR3DL2* alleles and some sequences extending 3’ past *KIR3DL2*. Figure 4 shows the alignment of the contigs relative to the reference haplotype MN167513 (cA01∼tB01) in the 18-probe experiment. It shows a small < 2 kb gap in the assembly in *KIR2DL3*. Otherwise, the contigs provide complete and overlapping coverage across the reference haplotype sequence. Supplementary Figure 1 contains the assemblies for all individuals in all three sets of experiments, along with Qualimap, NanoPlot, and Quast reports.

**Table 1.** Assembly statistics. For each of experiments in the rows, shown is the population, number of capture probes, the number of contigs spanning 75% of the haplotype (LG75), and the coverage of and concordance to the reference.

| pop. | probes<br># | LG75 | coverage<br>% | concordance<br>% |
| --- | --- | --- | --- | --- |
| EUR | 200 | 1.1 | 98% | 99.98% |
| EUR | 15 | 1.3 | 93% | 99.98% |
| AFA | 15 | 1.6 | 92% | 99.98% |
| **AFA** | **18** | **1.8** | **97%** | **99.97%** |

**Figure 3.**
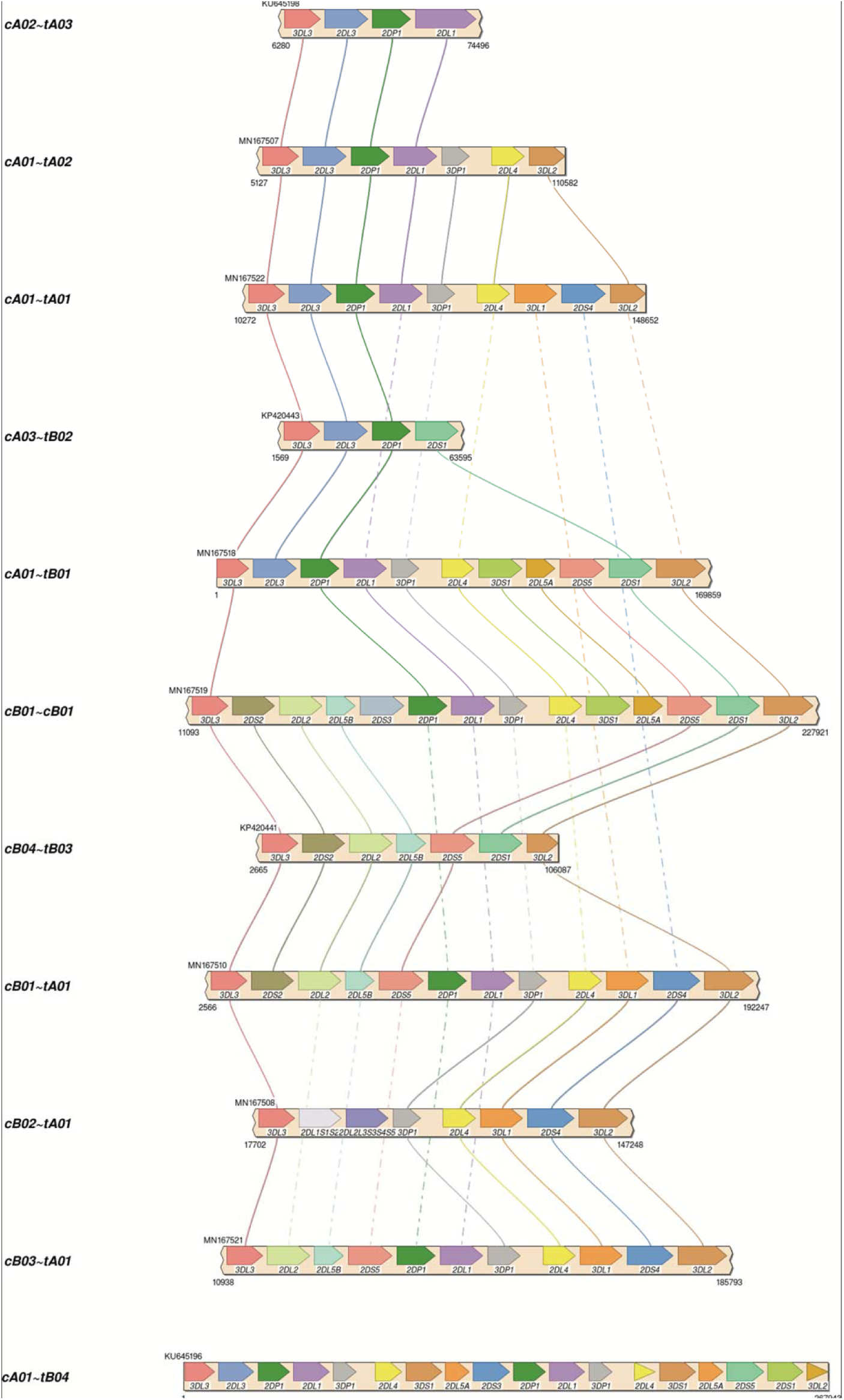
Haplotypes structures in the two validation cohorts. Each haplotype represents one of the structures in the two validation cohorts. The unofficial name of the haplotype is on the left. Lines connect genes with the same name in different structures. Solid lines connect the same gene in neighboring structures. Dashed lines connect the same gene in non-neighboring structures (i.e., the line goes through one or more neighboring haplotypes). cA02∼tA03: 1 EUR. cA01∼tA02: 1 AFA. cA01∼tA01: 5 AFA, 9 EUR. cA03∼tB02: 1 EUR. cA01∼tB01: 1 AFA, 1 EUR. cA01∼tB04: 1 EUR. cB01∼cB01: 2 AFA, 1 EUR. cB04∼tB03: 1 EUR. cB01∼tA01: 3 AFA. cB02∼tA01: 1 EUR. cB03∼tA01: 1 AFA. The visualization tool (SimpleSynteny) supports up to 10 sequence comparisons, s cA01∼tB04 is included at the bottom without explicit connections to other haplotypes.

**Figure 4.**
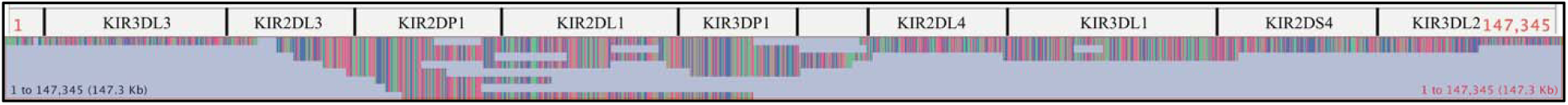
Alignment of assembled contigs with reference haplotype sequence MN167513 (cA01∼tA01), whose length is 147,345. The gene features are annotated across the top. The contigs are stacked below and colored by nucleotide.

The optimized 18 capture probe provided results very similar to the full candidate set of 200. Since this is the most efficient method, this probe coverage is further explained below. The 18 probes covered the haplotypes to an average distance of 2398 bases. Figure 5 shows how the probes are distributed across a typical 19 kb region. The image shows an alignment displayed in Integrative Genomics Viewer (IGV) of the set of 18 probes to cB01∼tB01 reference KP420442. The top of the image shows it is zoomed into 49 kb of the haplotype (∼50-100kb). In the middle track, the vertical ticks with the red numbers above indicate the alignment locations of the probes, with the red number being the label of the probe. In the bottom track, the horizontal blue lines indicate the locations of exons and repetitive elements. The probe locations avoid the blue variable (exons) and repetitive (Alus, LINEs, etc.) regions but achieve complete coverage to a resolution of less than 5 kb across the 49 kb. Only 7 distinct probes align to this region. From left to right, the probe sequence 4-3-12-10-2-7-13 occurs three times, except probe 10 does not align in the middle group. This alignment demonstrates how homology can be used to capture continuous KIR DNA over long distances with few probes without capturing off-KIR DNA. The set of 18 probes are included in Supplementary Table 2.

**Figure 5.**
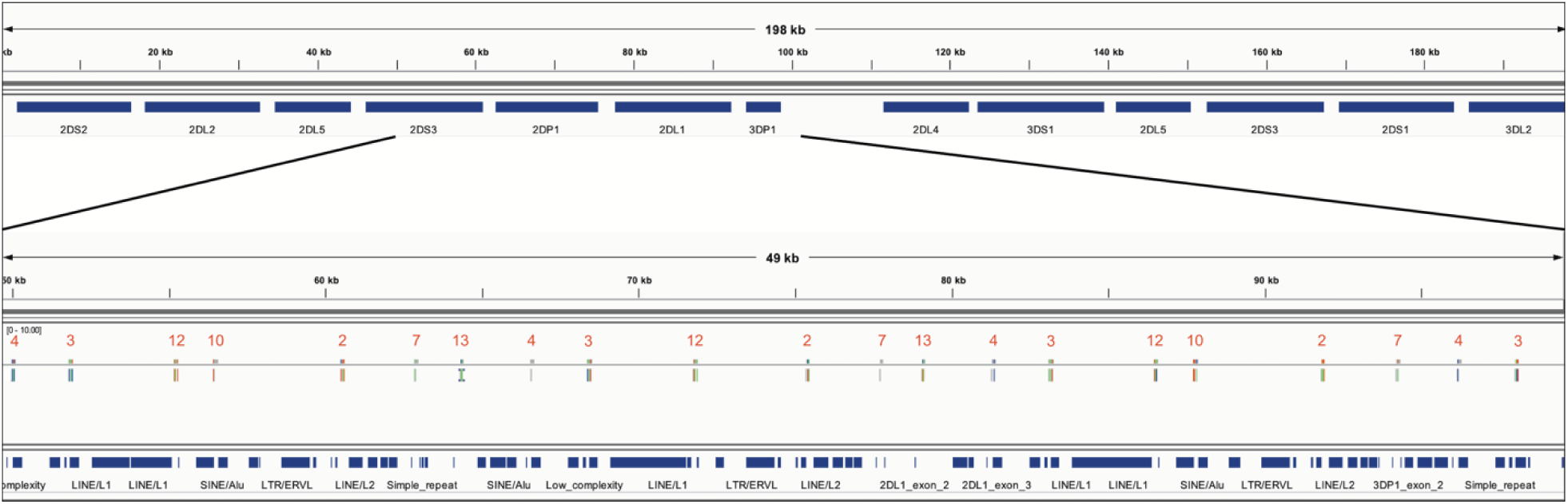
IGV depiction of the alignment of the 18 set capture probes across a 49 kb region of KP420440 (cB01∼tB01). The locations of the probes are displayed by the vertical ticks with the red labels above them in the middle track. The locations of exons and repeat elements (horizontal blue bars) are on the bottom track. 7 distinct probes align in this window. The probe pattern 4-3-12-10-2-7-13 repeats three times from left to right, except probe 10 does not align in the middle group.

The CCS reads in the 18-probe experiment provided an average of 47x coverage of all haplotypes, except for a small gap in all alleles of *KIR2DL2* and *KIR2DL3*, and a few alleles in other genes such as *KIR2DS2*. The gaps were ∼100 bases long, lead to gaps in the assembly < 2 kb, and were most likely introduced during PCR amplification of a repeat-rich region. See the reports in Supplementary Figure 1 for more information.

Using the 18 sequences as virtual probes, the WGS assembled into a paternal cA01∼tB01 (KIR3DL3*00101∼KIR2DL3*00101∼KIR2DP1*NEW∼KIR2DL1*00302∼KIR3DP1*0030202∼ KIR2DL4*0050101∼KIR3DS1*01301∼KIR2DL5A*00101∼KIR2DS5*00201∼KIR2DS1*00201 ∼KIR3DL2*00701) haplotype and a maternal cB05∼tB01 (KIR3DL3*00301∼KIR2DS2*005∼KIR2DP1*NEW∼KIR2DL1*0040105∼KIR3DP1*0030202∼ KIR2DL4*00501∼KIR3DS1*01301∼KIR2DL5*00101∼KIR2DS5*00201∼KIR2DS1*00201∼KI R3DL2*NEW); the assembly and its annotations are included in Supplementary Figure 2. The distal/telomeric halves are mostly homozygous, with approximately a dozen variants between them. This is the first occurrence in the genome databases of a cB05 with tB01 in the same haplotype; cB05 has *KIR2DS2*/*KIR2DS3* fusion (named KIR2DS2*005 in IPD-KIR). KPI(37) (preprint), an algorithm to genotype KIR from WGS, confirms cB05∼tB01 with cA01∼tB01 as the most-likely pair of structural haplotypes. The recently published PacBio full-genome assembly of this individual(36) assembled the cA01∼tB01 in paternal contig SRHB01000968.1, but the maternal contigs do not contain any KIR haplotypes.

The code to assemble and annotate KIR haplotypes from CCS reads, including an example, is located at https://github.com/droeatumn/kass. The ‘main’ workflow performs the assembly. The ‘annotate’ workflow labels the genes, exons, and introns in GenBank’s .gbf and .tbl formats. The ‘align’ workflow aligns the contigs to a reference and produces reports with which to evaluate the assembly. The code is supported by a Docker container at https://hub.docker.com/repository/docker/droeatumn/kass, for convenient execution. The minimum recommended hardware for targeted sequencing is 30G main memory and 8 CPU cores. More of each is helpful, especially with WGS. On an Ubuntu 18.04 Linux server with 40 core (Intel Xeon CPU E5-2470 v2 @ 2.40GHz) and 132G main memory, a single targeted assembly (∼70M fastq.gz) averaged 66 minutes and the WGS (∼70G fastq.gz) was 69 minutes. On MacOS 10.15.5 with 4 core (2.7 GHz Quad-Core Intel Core i7) and 16G main memory, a single targeted assembly averaged 125 minutes. Average times are reduced when assemblies are run in parallel.

## Discussion

These experiments in individuals from diverse populations demonstrate that KIR haplotypes can be efficiently enriched, sequenced, and assembled by using only 18 capture probes. The workflow successfully reconstructed all haplotypes from targeted sequencing in 16 individuals and WGS in 1 individual. Recent advances in Single-Molecule Real-Time sequencing by PacBio have improved quality and extended its applicability to WGS and highly repetitive regions. We have confirmed successful assembly with multiplexing up to eight individuals, which, we estimate, should lead to costs that rival full-exon short-read sequencing, and an order-of-magnitude more efficient than fosmid-based library preparation.

A 2016 manuscript(38) describes PING, which is software to interpret KIR from short (<=300 bp) reads. It uses probes to capture 800 bp DNA fragments. Although the total number of probes required for KIR capture was unspecified, the total number for KIR and HLA was 10,456. PING’s highest resolution results are obtained by aligning the short reads to full-gene references, calling the SNP variants, and then calling the two most likely reference alleles given the SNP genotypes. Although a great improvement over other current technologies at the time, the PING method doesn’t phase/link variants within a gene or the haplotype as our long-read sequencing and capture method allows. Further, PING uses more than an order of magnitude more capture probes, which can be expensive to capture shorter fragments. It produces probabilistically phased lower-resolution predictions compared with our long-read assembly, which produces linked multi-gene and haplotype sequences without references. Therefore, we feel our method appreciably adds to the ability to properly analyze KIR regions by leveraging long read technologies for increased resolution and phasing compared with the previous NGS approach and also by leveraging high-throughput library preparation for reduced cost compared with the previous haplotyping approach.

In addition to demonstrating an efficient targeted haplotyping strategy, to the best of our knowledge, this the first report of KIR full diploid haplotype assembly from WGS. Our approach was able to assemble both haplotypes from WGS whereas the previously reported whole-genome assembly could not, underlying the necessity of a KIR-specific assembler. Both regions comprising the haplotype it missed (cB05∼tB01) are not in the primary human genome reference, and the two have not been reported together previously. Perhaps this lack of representation in the reference contributed to the missing assembly. Other possibilities include the lack of binning/separation of KIR reads from the rest of the genome before assembling, or differences in the tools used in the workflow. Regardless, this experiment demonstrates the value added by the bioinformatics algorithms, in addition to the targeted capture and assembly.

The suspected amplification problem causing the small gap in *KIR2DL2L3* occurs in a ∼100 base region of poly-ATs, with L1s (and ALUs) on either side. It appears most or all PCR methods have a problem with this region, as almost every *KIR2DL2* and *KIR2DL3* reference allele has a different poly-AT sequence for this region (Supplementary Figure 3); these reference alleles were sequenced on various platforms but generally (if not fully) amplified with PCR. The PCR-less WGS from GIAB have no gaps. Since *KIR2DL2* or *KIR2DL3* occur in most haplotypes and occur ∼10-20% from the proximal end, their short gap limits LG75 to its reported value of 1.8 in the AFA cohort. In cases where the gap is not an issue, such as WGS, LG75 will probably be 1, and LG100 will probably be a better metric.

The power of this method to assemble repetitive KIR regions without incorporating false non-KIR genomic signals may lie in the strongest recombination hotspot, the 10 kb intervening region between *KIR3DP1* and *KIR2DL4*. Conventionally, KIR haplotype names (e.g., ‘cA01∼tA01’) have been described as two halves (‘c’ and ‘t’) separated by a recombination hotspot (‘∼’). The rate of recombination between the two halves is so frequent that any two may be found with each other, despite the relative evolutionary youth of the region. This hotspot stretches over 9 kb between *KIR3DP1* and *KIR2DL4*. Any two alleles of this region are over 99% identical and consist of 13% Alus (SINEs) and 58% LINE1 repeat elements. Figure 6 shows an alignment of the region to itself. The top part of the figure displays the location of the *KIR3DP1*-*KIR2DL4* intergenic region in the context of the cA01∼tA01 primary human genome reference. The bottom half zooms into the intergenic region and shows a dot plot of the alignment. The lines on the dot plot indicate stretches of the haplotype that align with itself, either in the same location or a different location. The red lines indicate matching in the same orientation as the overall haplotype, and the blue indicate matching in the reverse complement. The red and blue horizontal bars at the bottom of the figure detail the location of repetitive elements. The red from 0-2000 simply shows that this region aligns with itself. There is a stretch of 3 kb from ∼5200-8200 that reverse complement matches ∼400-3400. The yellow boxes highlight Alu repeats: AluSx3 from ∼800-1000 is matched in the reverse complement by AluSx1 (∼6400-6600) and AluSx4 (∼7100-7400) and the same orientation by AluSq2 (∼8200-8400). All elements are surrounded by very similar L1 elements for at least 1000 bp on both sides. This stretch of 3000+ bases of reverse complement repeats provides fertile ground for homologous recombination between the two halves of KIR haplotypes. This is the most difficult region to phase and the results demonstrate that the combination of variants and read length is generally high enough to phase full haplotypes with diploid reads. Although this region is an extreme example, the other recombination sites, which are usually internal to genes and result in gene fusions, homologously recombine via the same elements(39)(40)(17)(41)(42), although only the *KIR2DL4*-*KIR3DP1* intergenic region contains as many elements in both directions(11). Our assay is the only high-throughput method that allows analysis of all of these regions.

**Figure 6.**
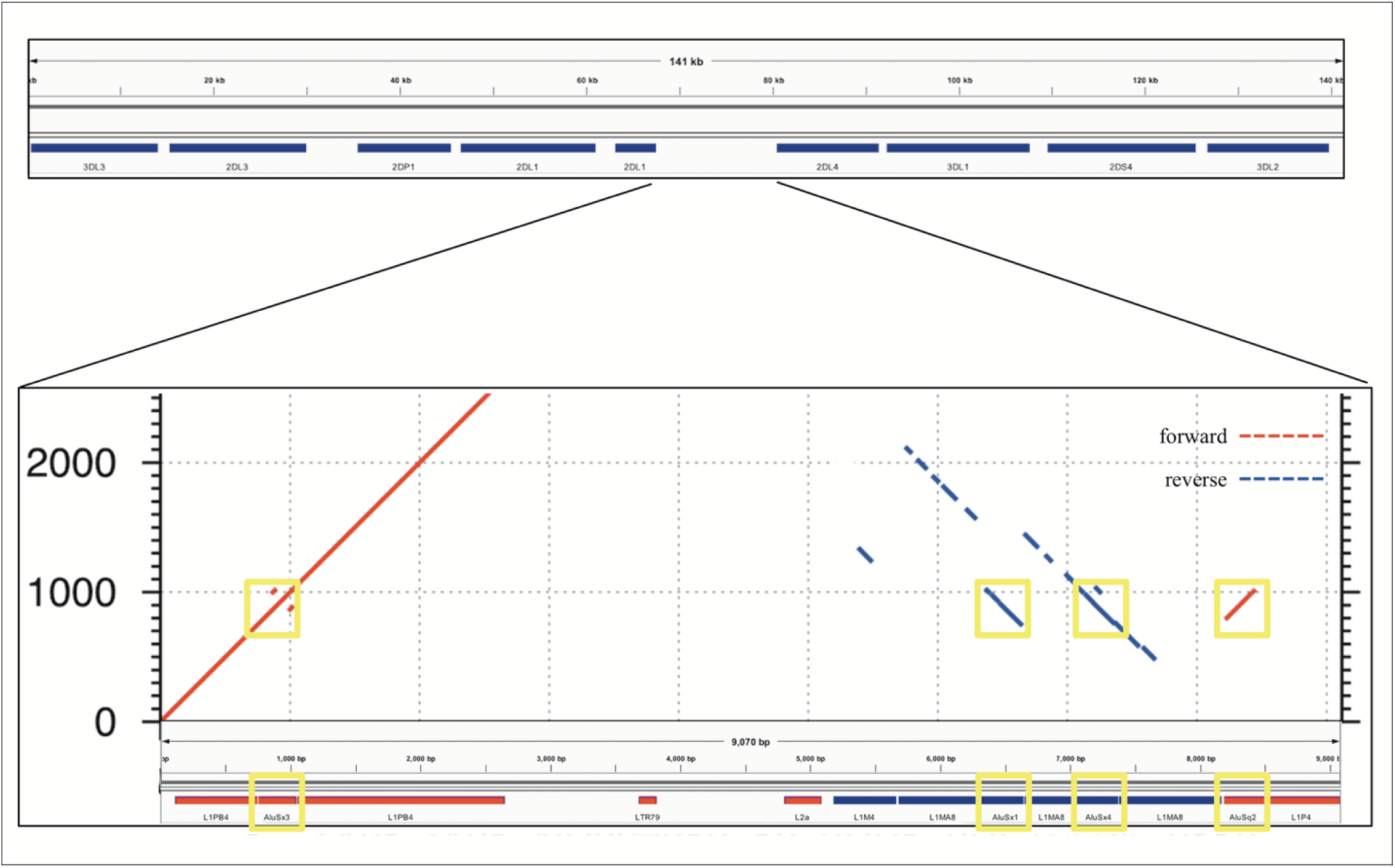
Recombination hotspot. The top shows the 9 kb haplotype context of the repetitive region of the *KIR3DP1*-*KIR2DL4* intergene region, whose self-alignment is shown in the bottom half. Red lines indicate alignment of the haplotype with itself in the same orientation, and blue lines indicate reverse complement orientation. The location of the repetitive elements is at the bottom, with red and blue again indicating orientation. The yellow boxes highlight three AluSx and one AluSq elements that align with each other, two in each direction.

Future efforts include expanding testing to other populations, resolving the *KIR2DL2L3* gap, expanding capture for some *KIR3DL2* alleles, expanding the assembly and annotation to bordering LILR genes, and optimizing multiplexing. Although the diverse AFA and EUR cohorts demonstrate proof of concept and expand our human genome references, it is important to develop reference sets for all populations. Expanding the capture would help ensure that *KIR3DL3* and *KIR3DL2* are sufficiently captured, help define any deleted haplotypes that may include these two genes and capture potentially relevant regulatory signals. Currently, all fully sequenced haplotypes contain some portion of these two bordering genes.

Our approach leverages sequence similarity across multiple loci that were created by duplication followed by variation. Since this this is a true for many gene families, our approach should be more generally applicable to other regions that have a mix of homologous and variable/repetitive regions relative fragment length and capture characteristics.

The application of KIR genetics in medical research such as immunity, reproduction, and transplantation is encouraging, but limited by the technical difficulties for high-resolution interpretations at large scale and low cost. Here, a KIR haplotyping workflow was presented that can provide full-sequence haplotypes at approximately the same cost as full exon or full gene. For the first time, it allows high-resolution KIR haplotypes in population-sized cohorts, as opposed to lower-resolution genotypes. The analysis pipeline uses domain knowledge to assemble reads generated via well-established sequencing techniques that is accurate enough for personalized precision medicine and scalable to populations. To this point, most KIR association studies focus on variation at only one locus or one functional class to associate, while keeping the rest of the haplotypes static. Future full-haplotype studies will help KIR researchers better study gene combinations, regulatory regions, recombination hotspots, self-regulation, and non-binding factors that influence disease phenotypes. This increased ability will provide completed sets of population-specific reference haplotypes which will, among other things, enhance imputation power of lower resolution data. It allows for new comparisons that will provide insight into evolution and make this region the best annotated in the human genome, despite its complexity. Lastly, this novel approach will provide the capability to discover genetic associations in medically relevant areas such as infections, transplantation, cancer susceptibility, autoimmune diseases, reproductive conditions, and immunotherapy.

### Ethics Statement

All subjects provided written informed consent for participation in research and the study, and consent were approved by the National Marrow Donor Program Institutional Review Board.

## Funding

Supported by a grant from the Department of the Navy, Office of Naval Research (N00014-19-1-2888).

## Acknowledgements

Thanks to Cynthia Vierra-Green, Julia Udell, and Richard Hall for advice and review.

## Supplementary Information

Supplementary Table 1. Cohort details. The first column contains the individual ID used for this study. The second column contains the GenBank accession number of the reference haplotype. Accessions that start with K are from the European cohort, and accessions that start with M are form the African American cohort. The third column contains the informal haplotype name.

Supplementary Table 2. Capture probes. The 18 capture probe sequences in fasta format.

Supplementary Figure 1. AFA and EUR contigs. A gzipped tar file containing the assembled contigs for all AFA and EUR assemblies. Also included are Qualimap, NanoPack, and QUAST reports.

Supplementary Figure 2. WGS contigs. A gzipped tar file containing the assembled contigs for maternal and paternal haplotypes for WGS assembly from GIAB individual NA24385.

Supplementary Figure 3. Multiple sequence alignment of reference KIR2DL2 alleles in IPD-KIR. The variation between columns 6706 and 6744 demonstrates extensive reported variation in a poly-AT region.

## Notes

### Competing Interest Statement

JW, KI, and JB are or were employees of Laboratory Corporation of America Holdings.
ND and CL are or were employees of Integrated DNA Technologies, Inc.

