## Supplemental Figure 1 for "Efficient Sequencing, Assembly, and Annotation of Human KIR Haplotypes": ccs999KIR7_18_8.contigs_MN167525_qualimap.pdf

### Qualimap Analysis Results

*BAM QC analysis*

*Generated by Qualimap v.2.2.1*

*2020/04/21 17:50:31*

### 1. Input data & parameters

#### 1.1. QualiMap command line

| |
|---|
| qualimap bamqc -bam ccs999KIR7_18_8.contigs_MN167525_sorted.bam -nw 400 -hm 3 |

#### 1.2. Alignment

| | |
|---|---|
| Command line: | bwa mem -k1800 -W9000 -r10 -A1 -B100 -O40 -E40 -L50 -t7 MN167525.fasta ccs999KIR7_18_8.contigs.fasta.gz |
| Draw chromosome limits: | no |
| Analyze overlapping paired-end reads: | no |
| Program: | bwa (0.7.17-r1188) |
| Analysis date: | Tue Apr 21 17:50:30 GMT 2020 |
| Size of a homopolymer: | 3 |
| Skip duplicate alignments: | no |
| Number of windows: | 400 |
| BAM file: | ccs999KIR7_18_8.contigs_MN167525_sorted.bam |

### 2. Summary

#### 2.1. Globals

| | |
|---|---|
| Reference size | 236,345 |
| Number of reads | 15 |
| Mapped reads | 15 / 100% |
| Supplementary alignments | 1 / 6.67% |
| Unmapped reads | 0 / 0% |
| Mapped paired reads | 0 / 0% |
| Read min/max/mean length | 2,152 / 123,929 / 32,562.13 |
| Duplicated reads (estimated) | 1 / 6.67% |
| Duplication rate | 7.14% |
| Clipped reads | 9 / 60% |

#### 2.2. ACGT Content

| | |
|---|---|
| Number/percentage of A's | 92,331 / 27.23% |
| Number/percentage of C's | 76,262 / 22.49% |
| Number/percentage of T's | 87,112 / 25.69% |
| Number/percentage of G's | 83,368 / 24.59% |
| Number/percentage of N's | 0 / 0% |
| GC Percentage | 47.08% |

#### 2.3. Coverage

| | |
|---|---|
| Mean | 1.4348 |
| Standard Deviation | 1.0566 |

#### 2.4. Mapping Quality

| | |
|---|---|
| Mean Mapping Quality | 47.85 |

#### 2.5. Mismatches and indels

| | |
|---|---|
| General error rate | 0.02% |
| Mismatches | 60 |
| Insertions | 11 |
| Mapped reads with at least one insertion | 33.33% |
| Deletions | 24 |
| Mapped reads with at least one deletion | 66.67% |
| Homopolymer indels | 91.43% |

#### 2.6. Chromosome stats

| **Name** | **Length** | **Mapped bases** | **Mean coverage** | **Standard deviation** |
|---|---|---|---|---|
| MN167525 | 236345 | 339101 | 1.4348 | 1.0566 |

### 3. Results : Coverage across reference

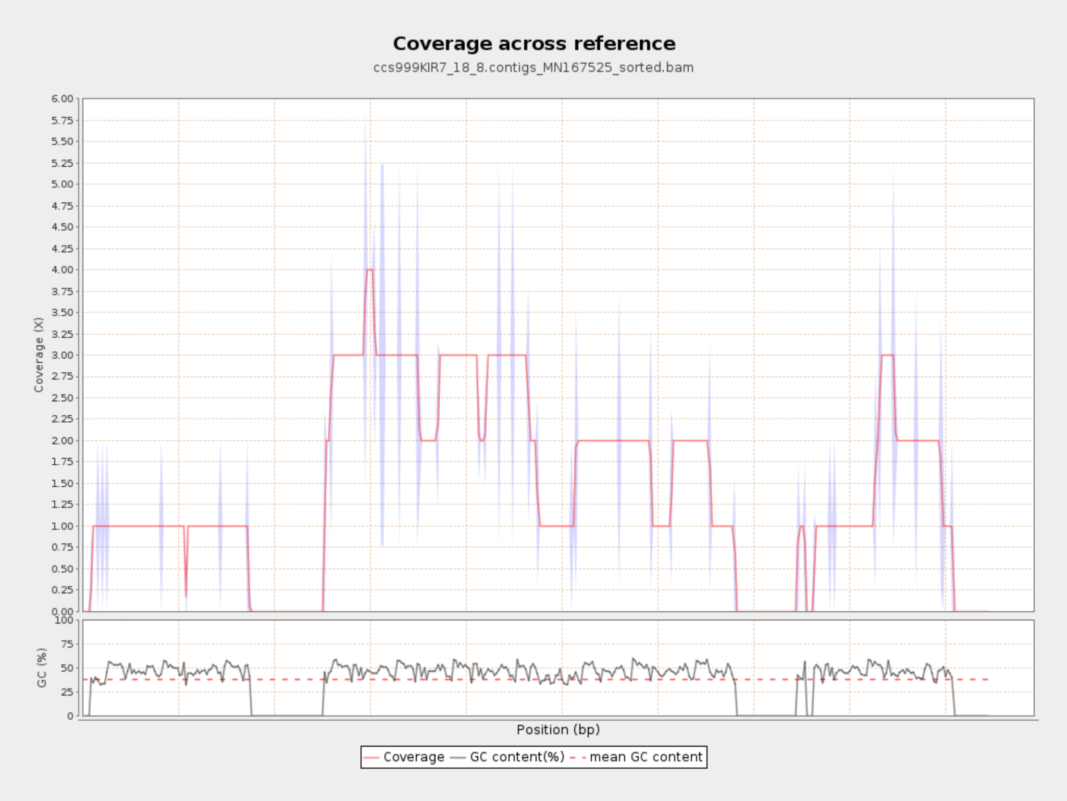

### 4. Results : Coverage Histogram

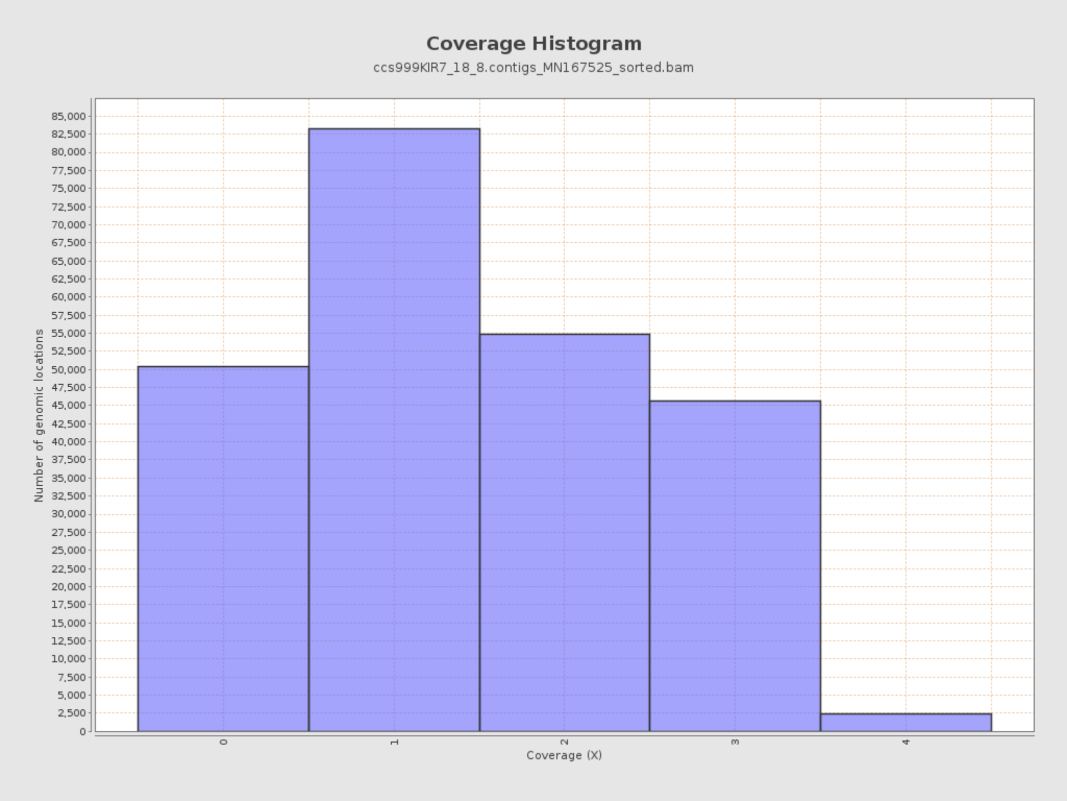

### 5. Results : Coverage Histogram (0-50X)

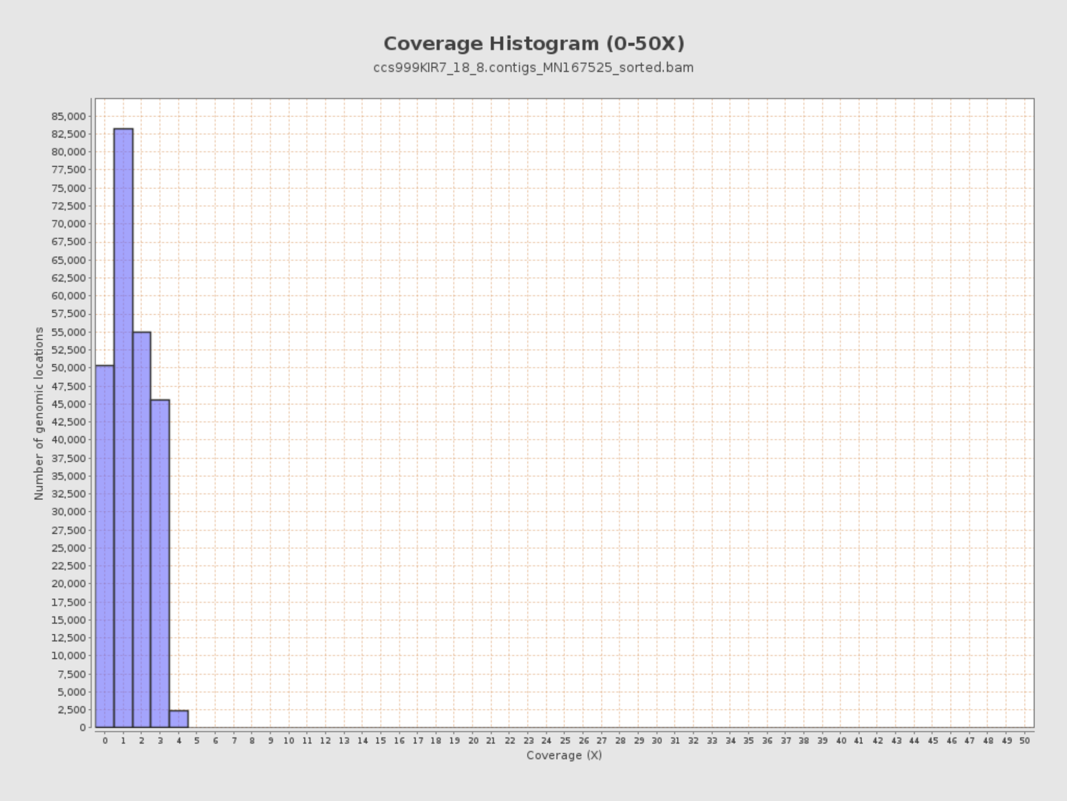

### 6. Results : Genome Fraction Coverage

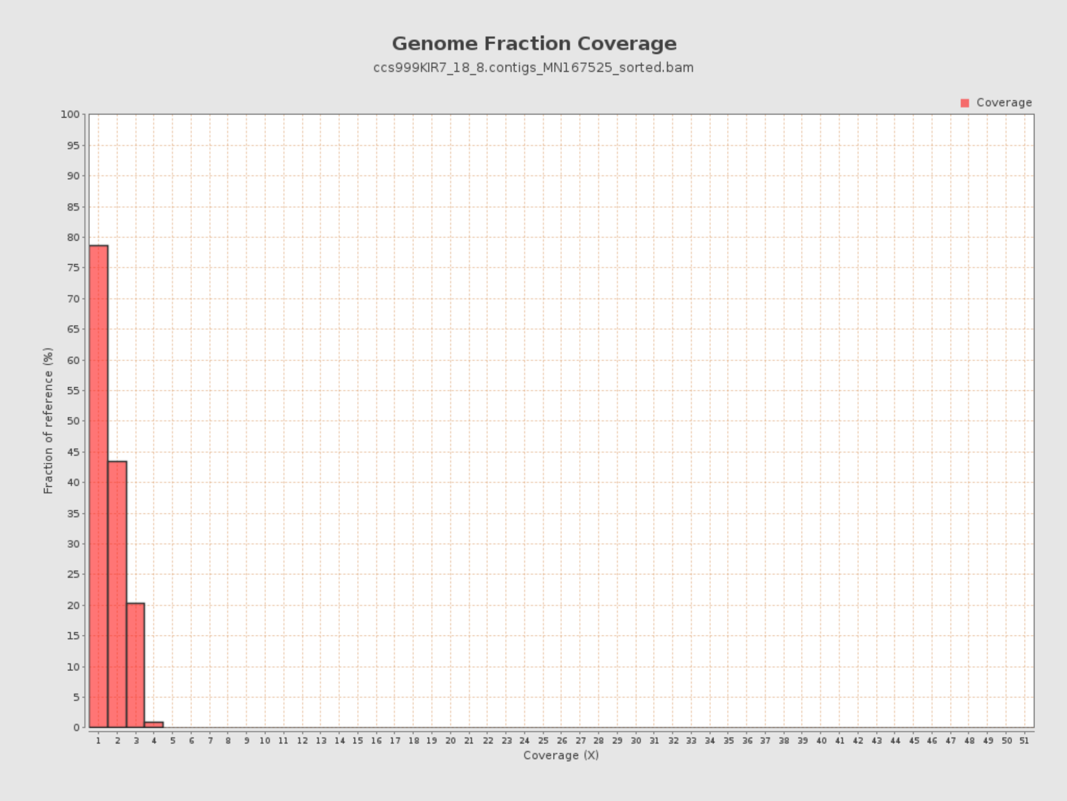

### 7. Results : Duplication Rate Histogram

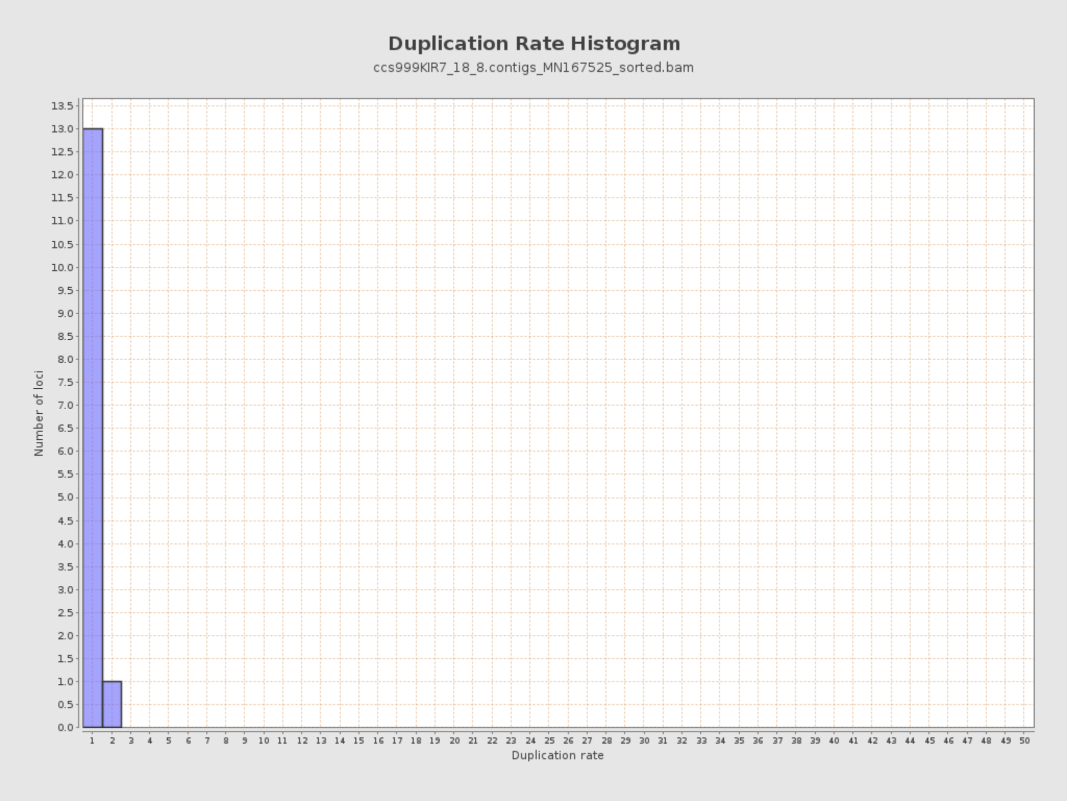

### 8. Results : Mapped Reads Nucleotide Content

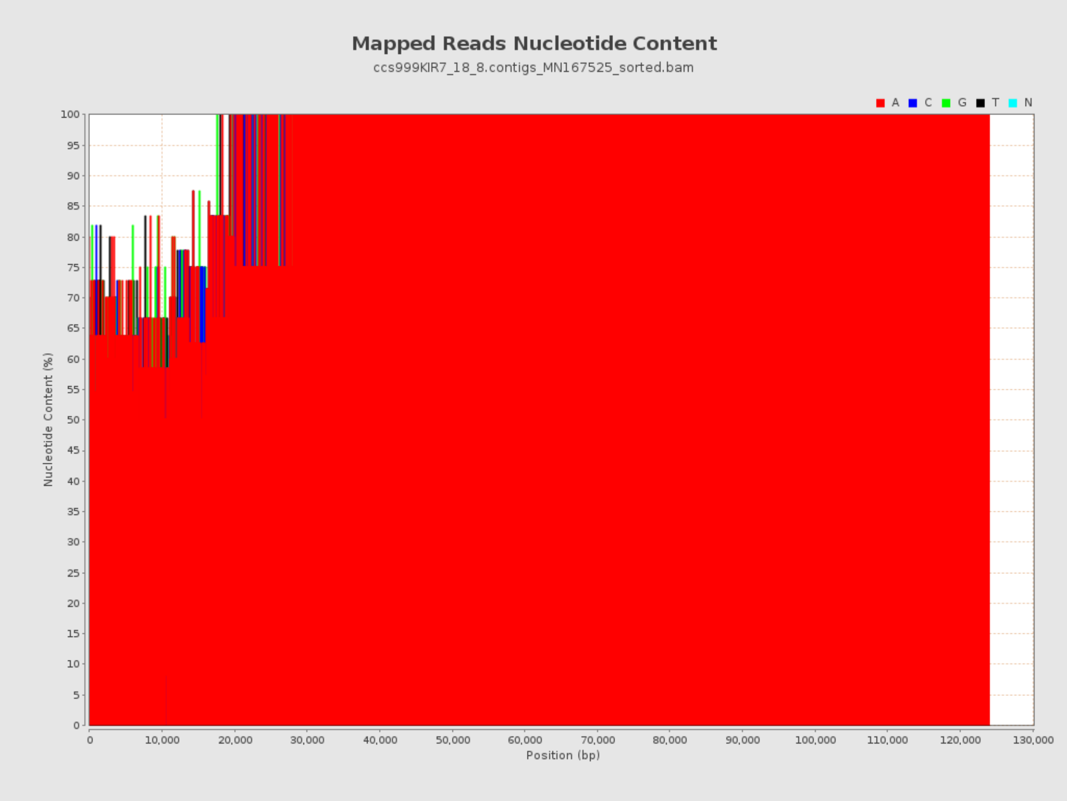

### 9. Results : Mapped Reads GC-content Distribution

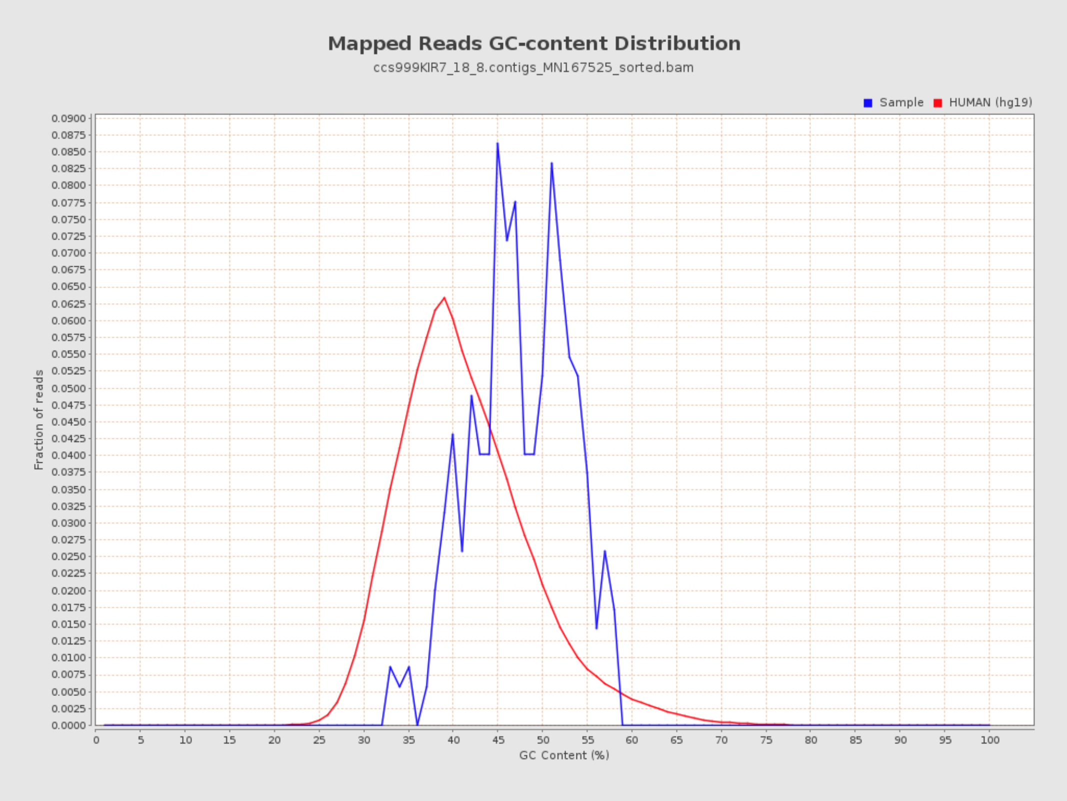

### 10. Results : Mapped Reads Clipping Profile

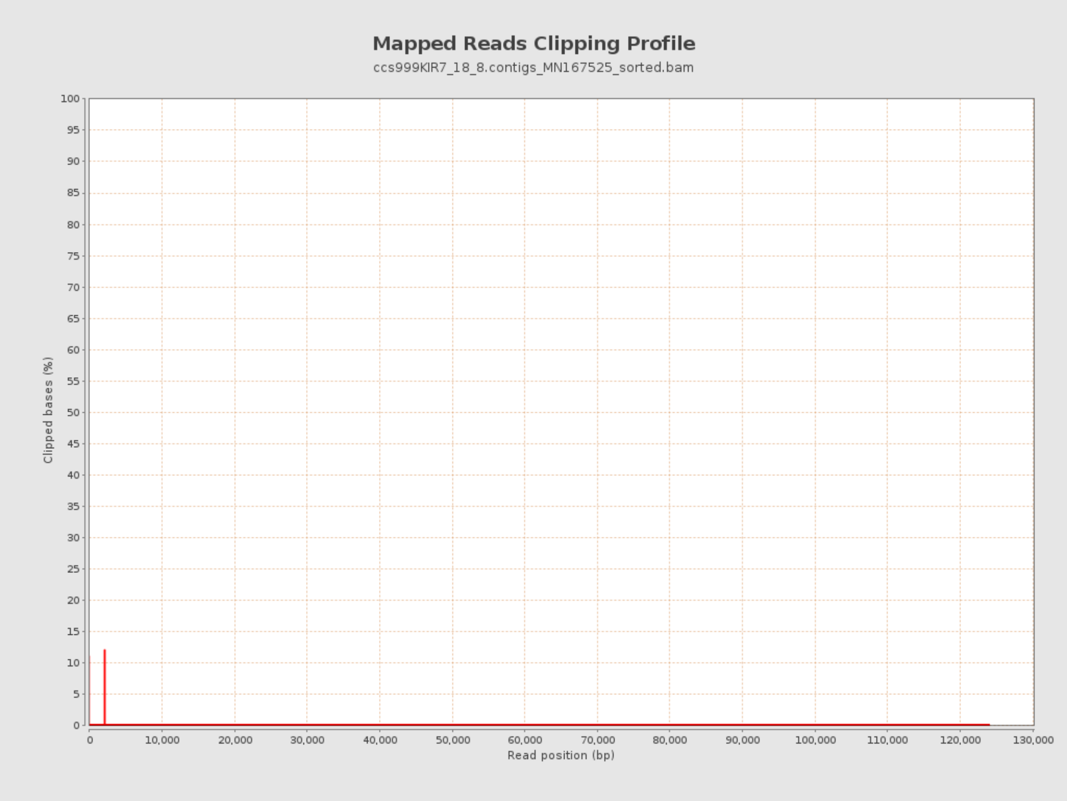

### 11. Results : Homopolymer Indels

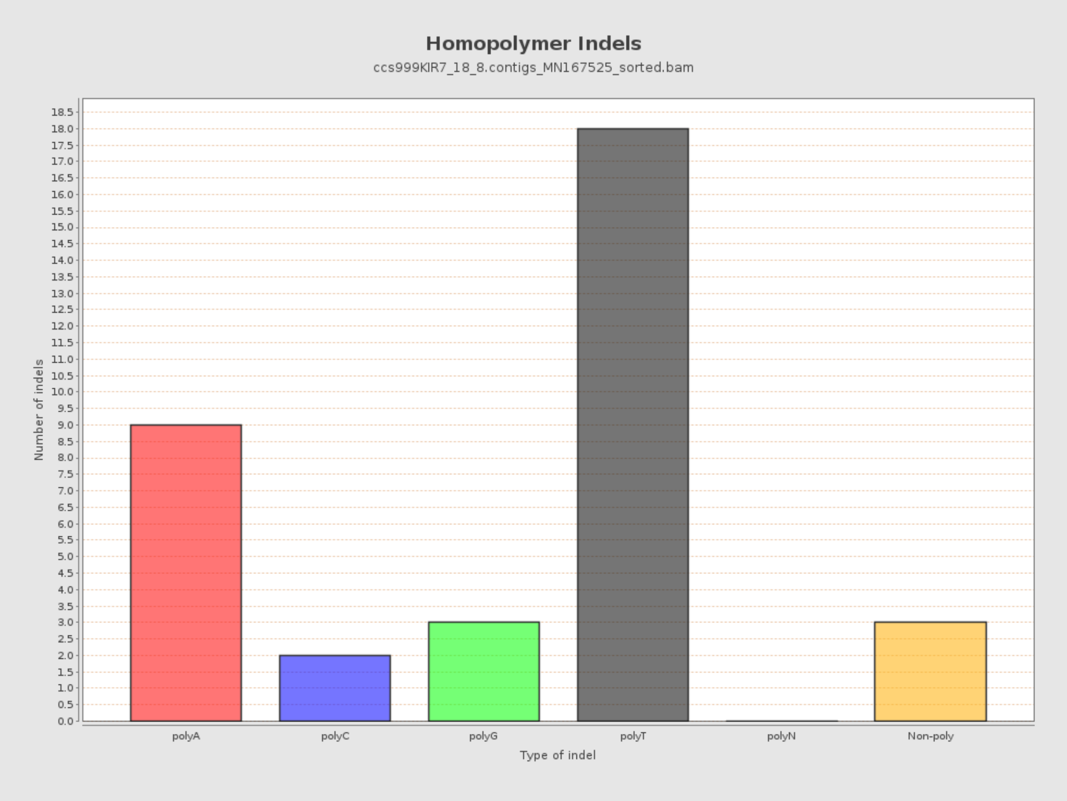

### 12. Results : Mapping Quality Across Reference

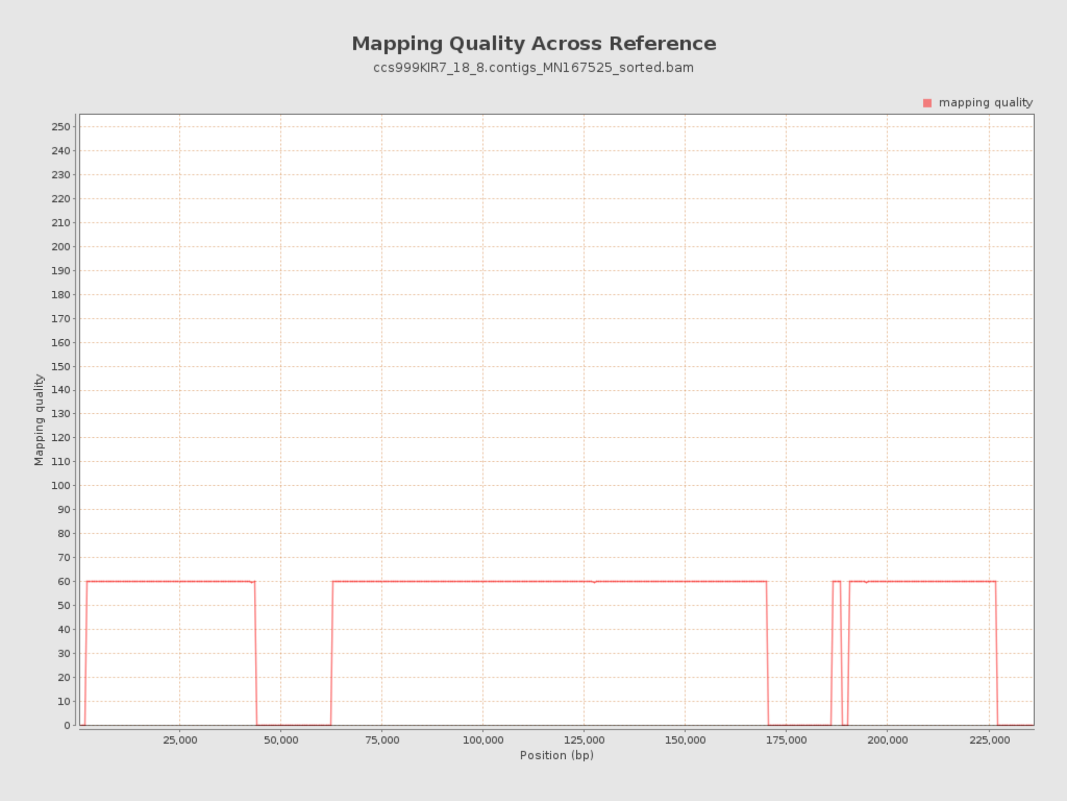

### 13. Results : Mapping Quality Histogram

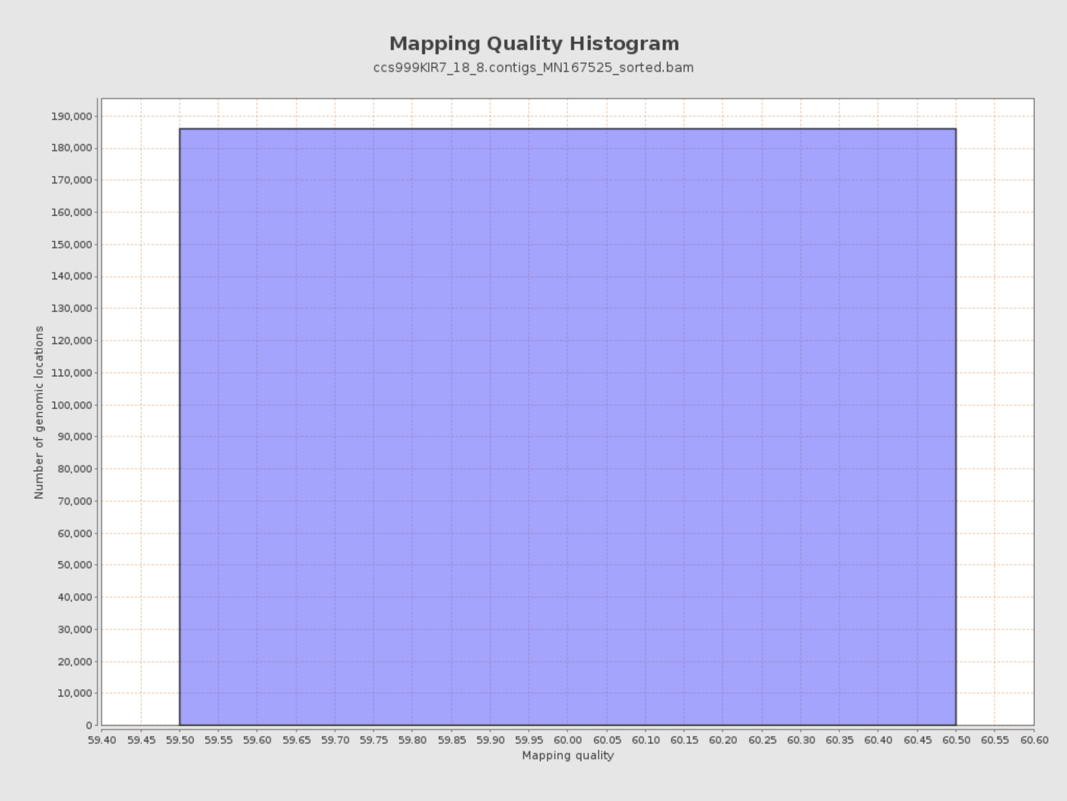
