## Supplemental Figure 1 for "Efficient Sequencing, Assembly, and Annotation of Human KIR Haplotypes": ccs999KIR7_18_8.contigs_MN167525_qualimap_genome_results.txt

BamQC report
-----------------------------------

>>>>>>> Input

     bam file = ccs999KIR7_18_8.contigs_MN167525_sorted.bam
     outfile = ccs999KIR7_18_8.contigs_MN167525_reports/genome_results.txt


>>>>>>> Reference

     number of bases = 236,345 bp
     number of contigs = 1


>>>>>>> Globals

     number of windows = 400

     number of reads = 15
     number of mapped reads = 15 (100%)
     number of supplementary alignments = 1 (6.67%)

     number of mapped bases = 339,101 bp
     number of sequenced bases = 339,073 bp
     number of aligned bases = 0 bp
     number of duplicated reads (estimated) = 1
     duplication rate = 7.14%


>>>>>>> Insert size

     mean insert size = 0
     std insert size = 0
     median insert size = 0


>>>>>>> Mapping quality

     mean mapping quality = 47.8452


>>>>>>> ACTG content

     number of A's = 92,331 bp (27.23%)
     number of C's = 76,262 bp (22.49%)
     number of T's = 87,112 bp (25.69%)
     number of G's = 83,368 bp (24.59%)
     number of N's = 0 bp (0%)

     GC percentage = 47.08%


>>>>>>> Mismatches and indels

    general error rate = 0.0002
    number of mismatches = 60
    number of insertions = 11
    mapped reads with insertion percentage = 33.33%
    number of deletions = 24
    mapped reads with deletion percentage = 66.67%
    homopolymer indels = 91.43%


>>>>>>> Coverage

     mean coverageData = 1.4348X
     std coverageData = 1.0566X

     There is a 78.71% of reference with a coverageData >= 1X
     There is a 43.5% of reference with a coverageData >= 2X
     There is a 20.27% of reference with a coverageData >= 3X
     There is a 0.99% of reference with a coverageData >= 4X
     There is a 0% of reference with a coverageData >= 5X
     There is a 0% of reference with a coverageData >= 6X
     There is a 0% of reference with a coverageData >= 7X
     There is a 0% of reference with a coverageData >= 8X
     There is a 0% of reference with a coverageData >= 9X
     There is a 0% of reference with a coverageData >= 10X
     There is a 0% of reference with a coverageData >= 11X
     There is a 0% of reference with a coverageData >= 12X
     There is a 0% of reference with a coverageData >= 13X
     There is a 0% of reference with a coverageData >= 14X
     There is a 0% of reference with a coverageData >= 15X
     There is a 0% of reference with a coverageData >= 16X
     There is a 0% of reference with a coverageData >= 17X
     There is a 0% of reference with a coverageData >= 18X
     There is a 0% of reference with a coverageData >= 19X
     There is a 0% of reference with a coverageData >= 20X
     There is a 0% of reference with a coverageData >= 21X
     There is a 0% of reference with a coverageData >= 22X
     There is a 0% of reference with a coverageData >= 23X
     There is a 0% of reference with a coverageData >= 24X
     There is a 0% of reference with a coverageData >= 25X
     There is a 0% of reference with a coverageData >= 26X
     There is a 0% of reference with a coverageData >= 27X
     There is a 0% of reference with a coverageData >= 28X
     There is a 0% of reference with a coverageData >= 29X
     There is a 0% of reference with a coverageData >= 30X
     There is a 0% of reference with a coverageData >= 31X
     There is a 0% of reference with a coverageData >= 32X
     There is a 0% of reference with a coverageData >= 33X
     There is a 0% of reference with a coverageData >= 34X
     There is a 0% of reference with a coverageData >= 35X
     There is a 0% of reference with a coverageData >= 36X
     There is a 0% of reference with a coverageData >= 37X
     There is a 0% of reference with a coverageData >= 38X
     There is a 0% of reference with a coverageData >= 39X
     There is a 0% of reference with a coverageData >= 40X
     There is a 0% of reference with a coverageData >= 41X
     There is a 0% of reference with a coverageData >= 42X
     There is a 0% of reference with a coverageData >= 43X
     There is a 0% of reference with a coverageData >= 44X
     There is a 0% of reference with a coverageData >= 45X
     There is a 0% of reference with a coverageData >= 46X
     There is a 0% of reference with a coverageData >= 47X
     There is a 0% of reference with a coverageData >= 48X
     There is a 0% of reference with a coverageData >= 49X
     There is a 0% of reference with a coverageData >= 50X
     There is a 0% of reference with a coverageData >= 51X


>>>>>>> Coverage per contig

	MN167525	236345	339101	1.4347712031140918	1.0566412411391957
