## Supplemental Figure 1 for "Efficient Sequencing, Assembly, and Annotation of Human KIR Haplotypes": ccs999KIR7_18_8.contigs_MN167525NanoPlot-report.html

NanoPlot Report

**Summary Statistics**

**Plots**

Histogram of read lengths

Histogram of read lengths after log transformation

Weighted Histogram of read lengths

Weighted Histogram of read lengths after log transformation

Dynamic histogram of Read length

Yield by length

Aligned read lengths vs Sequenced read length plot using dots

Aligned read lengths vs Sequenced read length plot using a kernel density estimation

Aligned read length vs Percent identity plot using dots

Aligned read length vs Percent identity plot using a kernel density estimation

Dynamic histogram of percent identity

### NanoPlot report

#### Summary statistics

| feature |  |
| --- | --- |
| General summary |  |
| Average percent identity | 100.0 |
| Mean read length | 32,562.1 |
| Median percent identity | 100.0 |
| Median read length | 19,934.0 |
| Number of reads | 15.0 |
| Read length N50 | 46,677.0 |
| Total bases | 488,432.0 |
| Total bases aligned | 339,085.0 |

#### Plots

##### Histogram of read lengths

  
  
  
  

##### Histogram of read lengths after log transformation

  
  
  
  

##### Weighted Histogram of read lengths

  
  
  
  

##### Weighted Histogram of read lengths after log transformation

  
  
  
  

##### Dynamic histogram of Read length

  
  
  
  

##### Yield by length

  
  
  
  

##### Aligned read lengths vs Sequenced read length plot using dots

  
  
  
  

##### Aligned read lengths vs Sequenced read length plot using a kernel density estimation

  
  
  
  

##### Aligned read length vs Percent identity plot using dots

  
  
  
  

##### Aligned read length vs Percent identity plot using a kernel density estimation

  
  
  
  

##### Dynamic histogram of percent identity
