## Supplementary figures and images for "Efficient Sequencing, Assembly, and Annotation of Human KIR Haplotypes"

### ccs999KIR7_18_8.contigs_MN167525PercentIdentityvsAlignedReadLength_dot.pdf

#### Aligned read length vs Percent identity plot

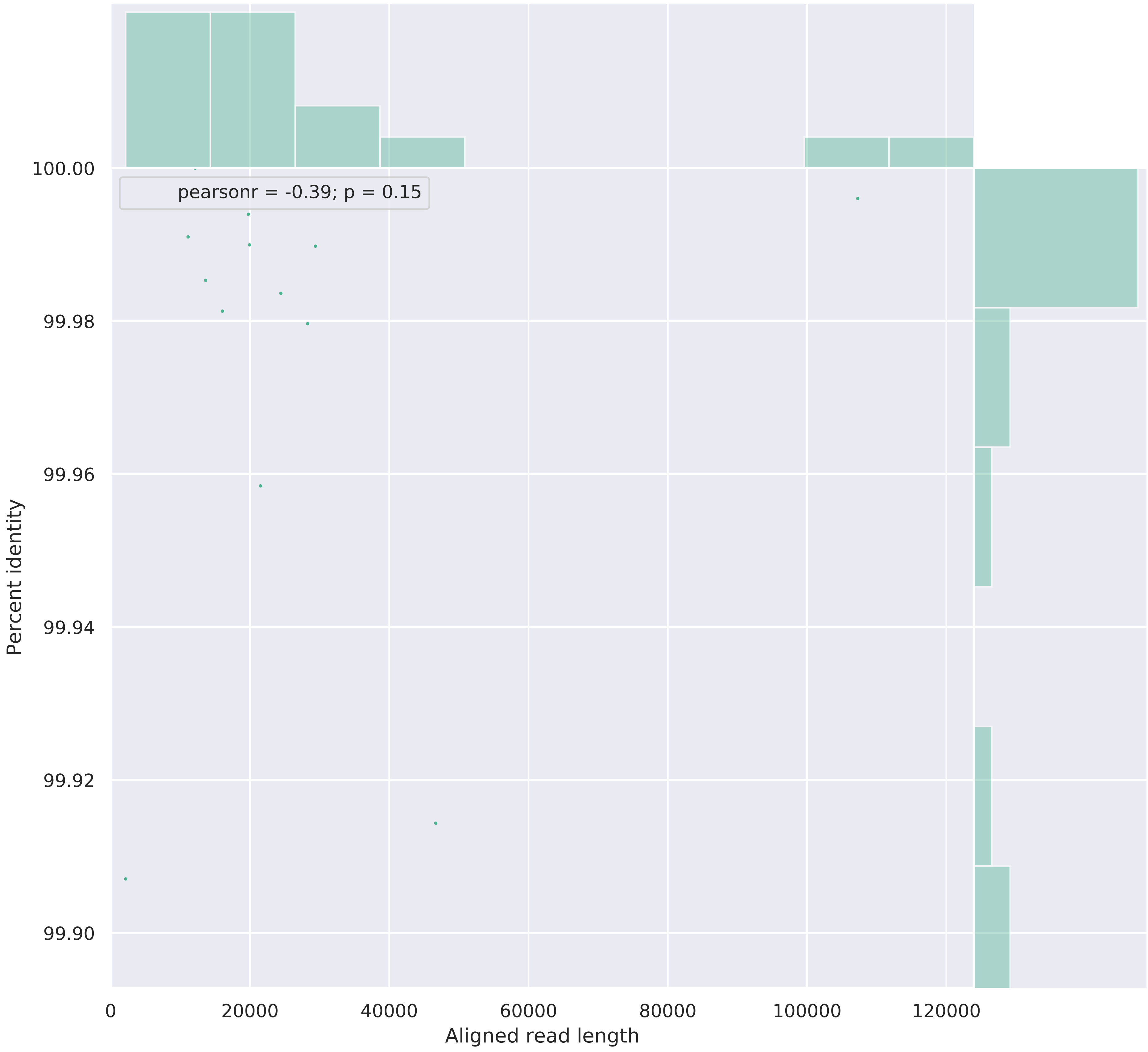

### ccs999KIR7_18_8.contigs_MN167525Weighted_HistogramReadlength.pdf

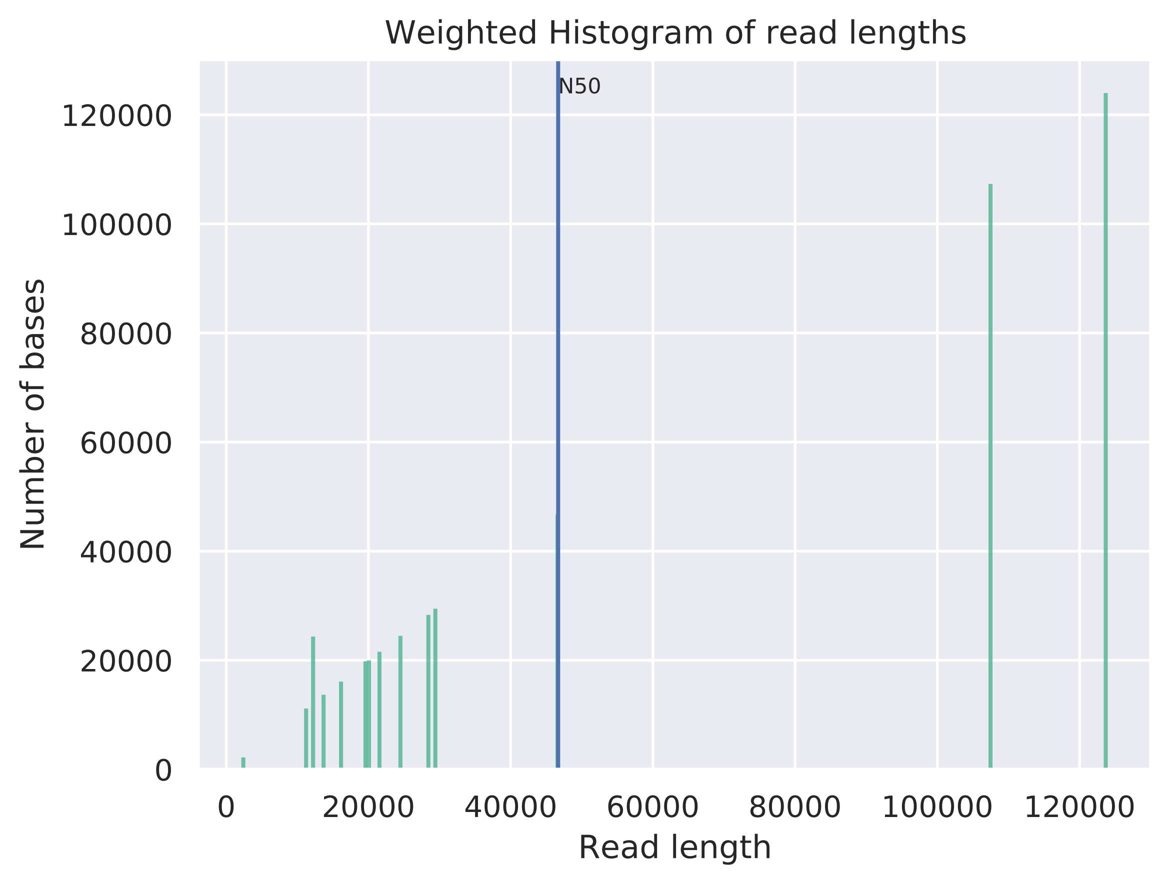

### ccs999KIR7_18_8.contigs_MN167525Weighted_LogTransformed_HistogramReadlength.pdf

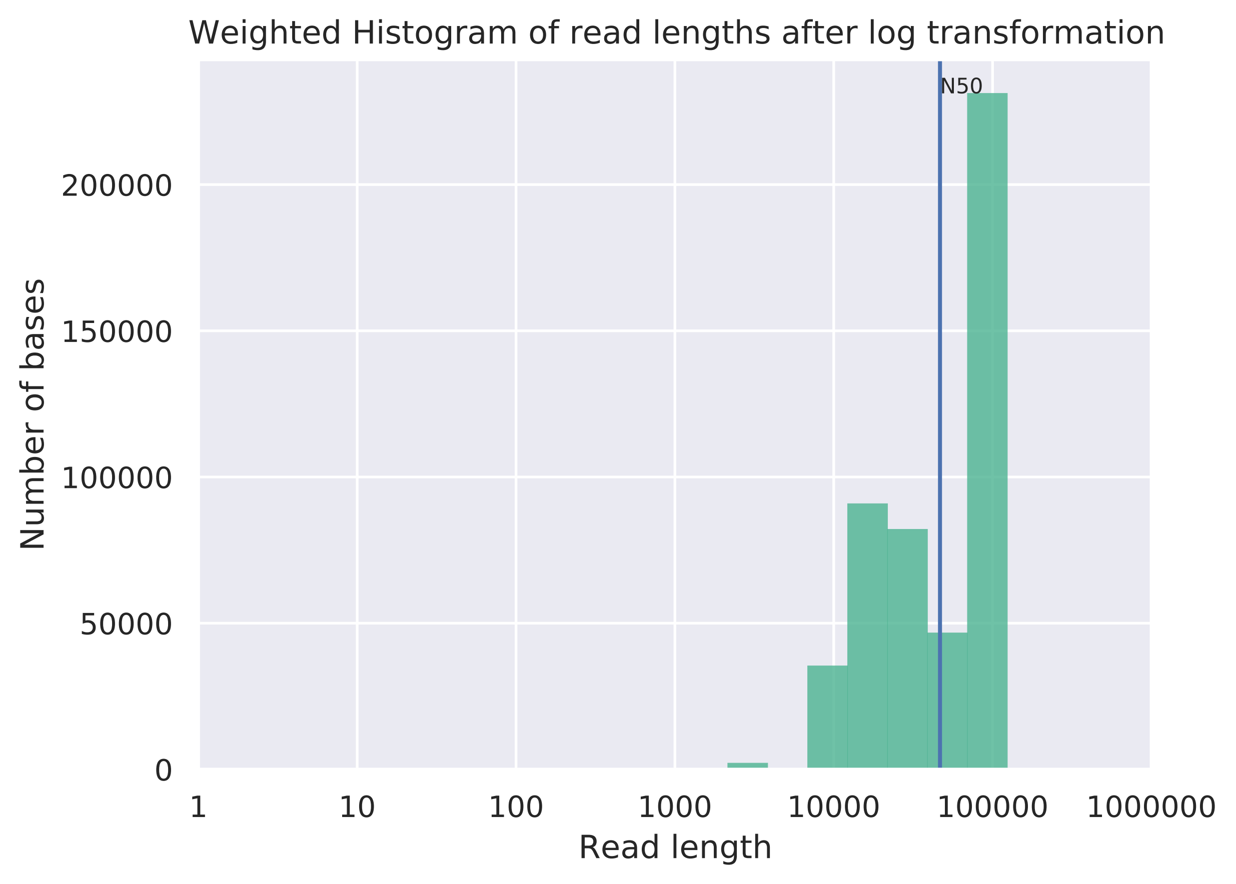

### Supplemental Figure 3

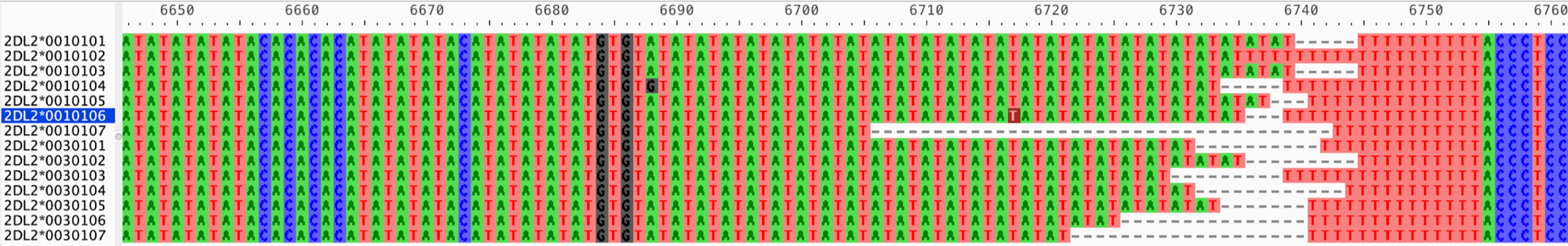
